# Intragenomic variants of a putative effector drive early-stage infection in a broad host-range rust fungus

**DOI:** 10.1101/2024.11.28.625939

**Authors:** Rebecca M. Degnan, Anne Sawyer, Donald M. Gardiner, Zhenyan Luo, Rebekah Frampton, Benjamin Schwessinger, Neena Mitter, Bernard J. Carroll, Grant R. Smith, Alistair R. McTaggart, Louise S. Shuey

## Abstract

Rust fungi are pathogens that impact plants of environmental, agricultural, cultural, and economic importance. Their mechanisms of pathogenicity are not well-understood but are likely governed by effectors, secreted proteins that manipulate host cellular processes to facilitate infection and suppress immune responses. We sought to understand how three effector candidates (*EFC1, EFC2*, and *EFC3*) expressed in the first stages of *Austropuccinia psidii* (myrtle rust) infection influence pathogenicity. We experimentally tested gene function through application of double-stranded RNA (dsRNA) and characterised the genomic landscape of putative effectors expressed during infection to assess whether putative effectors are needed for infection, and whether they are under selection pressure. One of the three screened candidates, *EFC1*, met our criteria of an effector in that it was predicted to be secreted, and was needed to cause but not maintain infection. We identified that this effector belongs to a gene family of intragenomic variants in tandem repeats flanked by transposable elements. Single nucleotide polymorphisms among these variants have signatures of non-neutral selection. This effector has predicted structural homology to a glycosaminoglycan-binding domain and may have a role in pectin or chitin-binding. We hypothesise that intragenomic variability in this family of effector genes facilitates host-range versatility in the *A. psidii*-Myrtaceae pathosystem.

## 1.0 Introduction

Rust fungi (Pucciniales) are plant pathogens that impact food security (Bettgenhaeuser et al., 2014), agricultural economies (Carretero et al., 2011), and natural environments, exemplified by invasive species that change landscapes and cause tree extinctions (Fensham and Radford-Smith, 2021; Pegg et al., 2018). Over 10,000 species of rust fungi are described and most are considered specific to a host species or genus of plants, to the extent that they are used as biological control agents against weeds in sensitive ecosystems (Wood, 2012), (Weidemann and Tebeest, 1990), (Gomez et al., 2008). Rust fungi that complete their life cycles on two hosts (heteroecious rust fungi), such as *Puccinia triticina* (the cause of wheat leaf rust), are considered specific to their two host genera (Bettgenhaeuser et al., 2014). Conversely, several species, including *Austropuccinia psidii* (myrtle rust), *Coleosporium solidaginis* (pine-aster rusts), and *Puccinia lagenophorae* (groundsel-ragwort rusts), are pathogenic to many hosts in their native or invasive ranges (Aime et al., 2018), (Müller-Schärer and Rieger, 1998). Myrtle rust is an extreme example, originating from Central and South America (Coutinho et al., 1998) but currently documented infecting at least 445 host species across 73 genera of Myrtaceae (Chock, 2020). Its spread to Australia impacted natural habitats and industry plantations of *Backhousia citriodora* (lemon-scented myrtle), caused decline in populations of endemic species, and threatens to cause the imminent extinction of at least 16 species of rainforest trees (Fensham and Radford-Smith, 2021; Pegg et al., 2017).

Many factors influence host specificity, including the landscape of proteins secreted by pathogens to maintain a compatible host-pathogen interaction. Obligate biotrophic fungi require a living host to survive and reproduce and maintain this interaction through establishment of haustoria (Voegele and Mendgen, 2003). Haustoria are established during the early stages of infection and facilitate uptake of nutrients from the host, and modulate trafficking of molecules, such as effectors, from pathogen to host. In the case of obligate biotrophic fungi, effector proteins are secreted either directly into the cytoplasm, the apoplast, or into the host cell via haustoria (Chaudhari et al., 2014).

Given the diversity of described rust fungi (over 7,000 species globally), relatively few effector proteins across a small number of species are functionally characterised (Ahmed et al., 2018; Aime and McTaggart, 2020; Prasad et al., 2019). This is partly due to the difficulty of working with an obligate biotrophic pathosystem, where traditional gene editing and other molecular techniques are often inviable (Petre et al., 2014). Foundational discoveries of effectors in rust fungi include Rust Transferred Protein (RTP1) in *Uromyces fabae* (bean rust), AvrM, AvrL567, AvrP4m and AvrP123 in *Melampsora lini* (flax rust), and PGTAUSPE10-1 in *Puccinia graminis* f. sp. *tritici* (wheat stem rust) (Ellis et al., 2007; Kemen et al., 2013, 2005; Upadhyaya et al., 2014b, 2014a). All six of these early-identified effectors are expressed in haustoria (Petre et al., 2014). RTPs were among the first fungal proteins shown to translocate from haustoria to host cells, and ultimately to the host nucleus (Kemen et al., 2005). RTP1, which inhibits cysteine proteases and promotes stability in the fungal extrahaustorial matrix through filament formation (Kemen et al., 2013; Pretsch et al., 2012), encompasses a multigene family conserved across the Pucciniales and found in ancestral species such as *Hemileia vastatrix* (coffee rust) (Lorrain et al., 2019).

Identification of candidate effectors in rust fungi has progressed through transcriptomics and genome-wide association studies, and while there has been significant progress in structure determination of wheat stem rust effectors, their functional characterisation remains a challenge (Lorrain et al., 2019; Outram et al., 2024). Ramachandran *et al*. (2016) used transcriptomics to identify 9 new secreted proteins (Shr1-9) from *Puccinia graminis* and *Puccinia striiformis*. Shr1-9 were characterised through a hypersensitive response (HR) suppression assay in *Nicotiana benthamiana*, and impacted multiple plant defence mechanisms (Ramachandran et al., 2017). Knowledge of the *P. striiformis* effector landscape has significantly advanced in the last 10 years with the molecular characterisation of numerous effectors including PSTha5a23, PstGSRE1, Pst18363 and Pst_8713, which suppress plant defences, disrupt ROS-induced host defences, and enhance *Pst* virulence (Cheng et al., 2017; Liu et al., 2022; Qi et al., 2019; Yang et al., 2020; Zhao et al., 2018). In the *Melampsora larici-populina* pathosystem, characterised effectors include CTP1–3, which contain N-terminal transit peptides that enable the proteins to translocate to the chloroplast stroma, and Mlp124478, which binds to plant TGA1a promotors. Effectors expressed by *Melampsora larici-populina* improve targeted trafficking within plant cells by mimicking plant specific sorting signals (Petre et al., 2016), repress immune response machinery in the host, and alter leaf morphology (Ahmed et al., 2018).

Two studies have identified putative effectors in *A. psidii*. Hayashibara *et al*. (2023) identified a putative effector in the MF-1 (Brazilian) strain of *A. psidii* that appears to localise to the nucleus but has not yet been functionally characterised *in planta* (Hayashibara et al., 2023). Frampton *et al*. (2024) identified several candidate effectors in the pandemic strain of *A. psidii* based on differential expression early in infection (Frampton et al., 2024). In controlled *A. psidii* inoculations, mitotic urediniospores germinate between 0–12 hours after inoculation on a leaf surface. Germ tubes differentiate into appressoria that apply pressure through an infection peg to directly penetrate the leaf cuticle. Up to 12 – 48 hours post-infection, the fungus develops a network of intercellular hyphae and haustoria within and around host cells. As disease symptoms become visible at six days post-infection and progress, *A. psidii* develops sori that contain thousands of new urediniospores that eventually rupture the leaf cuticle, creating a new lesion and restarting the infection cycle (Degnan et al., 2023b).

A significant gap in understanding is how rust fungi manipulate their hosts at early stages of infection, before the development of haustoria. We aimed to characterise three of the early-expressed candidate effectors identified by Frampton *et al*. (2024) to understand their role in the infection process (Frampton et al., 2024). We broadly hypothesised that these genes are crucial for early infection, but not to maintain an ongoing infection. We tested specific hypotheses that (i) infection by *A. psidii* would be supressed *in planta*, but *in vitro* germination would not be impacted, when targeting putative effectors with sequence-specific dsRNA, and (ii) infection by *A. psidii* would not be lessened by targeting these genes with curative dsRNA treatments at 3 days post-infection. One of the putative effectors, *EFC1*, was a gene family of intragenomic variants, and we hypothesised that duplicated effector variants may confer an evolutionary advantage and be maintained under selection. We use *in silico* analysis to describe the extent of genetic variation in the *EFC1* gene family, as well as previously developed dsRNA-based *in vitro* and *in planta* co-inoculation and curative assays to define the impact that members of the gene family have on the infection process (Degnan et al., 2023b, 2023a). The effector landscape of *A. psidii* may provide answers to the broad host range of *A. psidii*. Further, this study expands our knowledge of effectors in obligate biotrophic pathogens and potentially provides another target to improve management outcomes for rust fungi.

## 2.0 Experimental Methods

### Austropuccinia psidii inoculum preparation and spore germination conditions

*Austropuccinia psidii* urediniospores were harvested from myrtle-rust infected trees in the field. Uredinospores were desiccated for 48 hours and sieved to separate from dirt or plant matter before being stored long-term in cryo-safe tubes at -80 ºC. To prepare inoculum, urediniospores were resuspended in sterile distilled water and 0.05% Tween 20 for a minimum of 30 minutes before use, to allow spores to rehydrate. Inoculum concentration was determined using a haemocytometer and adjusted to 1×10^6^ spores/mL. In both detached leaf and *in vitro* assays, *A. psidii* urediniospores were germinated for 24 hours in a controlled environment room at 18ºC, 75% relative humidity, and with no lights.

### Plant propagation and preparation

*Syzygium jambos* plants, used in detached leaf assays, were clonally propagated from cuttings and grown under glasshouse conditions (28ºC, natural light, hand-watered once per day) at the Ecosciences Precinct, Dutton Park for 3 years. Over these years, plants were cut back and fertilised at regular intervals to encourage production of new growth. Only new, emerging growth was used in detached leaf assays.

### Selection of target genes and primer design

Putative effectors *EFC1* (*APSI_P014*.*1260*.*t1*), *EFC2* (*APSI_P001*.*5292*.*t1*), and *EFC3* (*APSI_P005*.*10948*.*t1*) were previously identified from a differential expression analysis in uninfected and myrtle rust-infected *Leptospermum scoparium* (Frampton et al., 2024). Primers were designed in Geneious Prime. At the 5’ end of forward and reverse primers, a T7 promotor sequence (5’–TAATACGACTCACTATAG–3’) was added to allow for transcription of sense and antisense strands in a single RNA synthesis reaction, resulting in dsRNA (Supplementary Table 1, Supplementary Figure 1).

### Cloning and identification of EFC1 divergent alleles

Phusion DNA polymerase-amplified *EFC1* PCR products were blunt-end cloned using the Thermo Scientific CloneJET PCR Cloning Kit, according to the manufacturer’s instructions. PCR products were ligated, and competent *Escherichia coli* cells were transformed with products via heat-shock and plated onto LB Agar Ampicillin-100. Plates were grown overnight at 37 ºC and 10 colonies were selected from each treatment for screening. Colonies were screened for the presence of the *EFC1* DNA insert using colony PCR with sequence-specific primers as described in the manufacturer’s instructions. Screened colonies that appeared to have been successfully transformed with the DNA insert, based on gel electrophoresis, were Sanger sequenced (AGRF) to determine the presence and number of divergent *EFC1* alleles.

### In vitro dsRNA synthesis

*A. psidii* urediniospores were manually homogenised in liquid nitrogen and RNA was extracted with TRIzol™ (ThermoFisher Scientific) according to the manufacturers protocol. cDNA to be used for PCR was synthesised from total *A. psidii* RNA with a ProtoScript® II First Strand cDNA Synthesis Kit, using the standard protocol (E6560, New England BioLabs). Target sequences were amplified with Phusion High Fidelity DNA Polymerase (ThermoFisher Scientific), under standard thermocycling conditions with recommended annealing temperatures.

dsRNA was synthesised, with T7 PCR products as templates, using the HiScribe™ T7 High Yield RNA Synthesis Kit (New England Biolabs). *In vitro*-synthesised dsRNA was treated with DNAse I and purified using a modified TRIzol™ (ThermoFisher Scientific) extraction, where precipitation was increased to 2 hours at -20ºC and RNA pellets were washed in 70% EtOH for 45 minutes to ensure complete removal of phenol contamination. Quality and concentration of dsRNA was determined with a combination of BioDrop analysis and gel electrophoresis.

### Urediniospore germination assays on polydimethylsiloxane discs

*A. psidii* spore inoculum (prepared as above) was combined via pipetting with dsRNAs at a concentration of 100 ng/μL and an aliquot of 20 μL was pipetted in 2 μL droplets onto each polyvinylpyrrolidone-coated polydimethylsiloxane (PVP-coated PDMS) disc (Soffe et al., 2019). Discs were incubated for 24 hours at 18ºC and 75% relative humidity to promote spore germination. Germination analysis of spores (percent germinated) was completed under a light microscope.

#### In planta assays

Two-to-three-year-old *S. jambos* trees were grown under glasshouse conditions and watered by hand once daily. To prepare for infection, trees were fertilised with Nitrosol at the recommended dose two weeks prior to infection. Trees were not fertilised while the assay was in progress. New growth on trees was sprayed adaxially and abaxially with approximately 1 mL per node of *A. psidii* inoculum (prepared as above) and either nuclease free H_2_O (negative control), or 100 ng/μL dsRNA (β*-TUB, GFP, EFC1, EFC2, EFC3, EFC1-01 (MK676_005456)* or *EFC1-19 (MK676_005467)*, depending on assay and treatment). Following inoculation, plants were incubated for 24 hours at 18ºC and 75% relative humidity, before being returned to the glasshouse. After 14 days, plants were photographed and either leaves were detached and scanned for pustule counting, or disease incidence (scored as percent coverage of diseased tissue) was assessed using the Leaf Doctor application^1^.

#### Phylogenetics, nucleotide diversity, and substitution rate

All cloned *EFC1* variants, as well as homologs in available *A. psidii* genomes, were aligned with MAFFT v7.490 and relationships were visualised with SplitsTree v4.14.8 (Katoh and Standley, 2013; Kloepper and Huson, 2008). Three sequences (*EFC1-26, EFC1-28* and *EFC1-06*) with early terminations (stop codons) were removed from the alignment used to produce sequence logo, pi plot, and dN/dS graphs. Logo size was 40×5 cm per line, with up to 100 symbols per line. The conservation of amino acids across variants was visualised using a sequence logo diagram prepared with WebLogo 3.7.5 (Crooks et al., 2004). The image was specified as 96 pixels/inch (dpi), with a boxed shrink factor of 0.5 and an x-axis height of 2 bits. We visualised nucleotide diversity as a pi plot, by calling SNPs across the codon alignment using SNP-tools (Page et al., 2016) and used vcftools (Danecek et al., 2011) to calculate pi (command --site-pi). Pi was plotted across the alignment using the ggplot2 package in R Studio (R Core Team, 2020).

Non-synonymous to synonymous substitution rate was used to evaluate the positive selection sites on the effector genes. The Multiple Sequence Alignment (MSA) and gene tree was used as the input for CodeML in PAML v4.9j (Yang, 2007). To run the site model, the following parameters were used: clock = 0, cleandata = 0, CodonFreq = 2, runmode = 0, NSsites = 0, 1, 2, 7, 8. Likelihood ratio tests (LRTs) with two degrees of freedom were performed to compare M1a-M2a and M8-M7. The Bayes Empirical Bayes (BEB) approach was used to calculate the posterior probability per positively selected site, which fit the condition ω > 1 and *P* > 0.95 (Deely and Lindley, 1981). To calculate rates of synonymous versus non-synonymous mutations, amino acid sequences of putative effector *EFC1* were aligned with MAFFT v7.490 (Katoh and Standley, 2013). PAL2NAL was applied to generate the corresponding protein-based codon alignment (Suyama et al., 2006). JC+I was selected as the best-fit model by ModelFinder and the gene tree was interfaced with IQTREE2 v2.0.7 with 10,000 ultrafast bootstraps (Hoang et al., 2018; Kalyaanamoorthy et al., 2017; Minh et al., 2013).

To identify orthologues of putative effectors in other rust fungi, tblastn was used to search publicly available genomes of rust fungi for homologs of all putative effectors. Potential targets were annotated with UniProt and aligned amino acids with Clustal Omega in Geneious Prime. IQtree was used to search for a maximum likelihood tree with a model test and 10,000 ultrafast bootstraps and 10,000 approximate likelihood ratio tests.

#### Genomic mapping

Sequences MK676_005453, MK676_005456 (*EFC1-01*), MK676_005467 (*EFC1-19*) and MK675_004127 were obtained with blastp 2.12.0+ (Johnson et al., 2008). Genomic positions of genes were determined with the corresponding GFF files, and the gene map was generated using the gggenomes packages (version 0.9.5.9000) (Hackl, Thomas et al., 2024).

#### Structural prediction and alignment

The structure of *EFC1-01* was predicted using the Alphafold 3 server, with the protein sequence as input (Abramson et al., 2024). To find homologous structures in the PDB, the *EFC1-01* protein sequence was searched in DALI (Holm, 2022). The closest match, determined by Z-scores and structural alignments, was superimposed with the Alphafold 3 *EFC1-01* prediction in PyMOL v1.3r1 (Schrödinger LLC, 2010). The isoelectric point and net charge of *EFC1-01* were calculated with Prot pi v2.2.

#### Small RNA (21-mer) prediction

A sliding window analysis was computed using the Biostrings package in R Studio 2023 to identify how many 21-base pair fragments would be identical between aligned *EFC1-01* and *EFC1-19* if they were diced into small RNAs (R Core Team, 2020). Data were plotted using ggplot2 and the rainfall_plot function (Wickham, 2020).

#### Statistics and reproducibility

For *in planta* assays, three biological replicates (trees) were used per treatment, with four technical replicates (newest leaves at top two nodes) per biological replicate. For *in vitro* assays, three biological replicates (PDMS discs) were used per treatment, with at least 100 urediniospores assessed per replicate. Technical replicates were combined to produce a single data point (biological replicate) for all statistical analyses and data visualisation. Welch’s two sample, two-tailed *t* tests were calculated at a 95% confidence interval using the dplyr package in R Studio 2024 (Hadley Whickam, 2021; R Core Team, 2020). *P* values of <0.05 were considered to be significant. Data were plotted using geom_box and geom_scatter functions within the ggplot2 package in R Studio 2024.04.1+748 and final aesthetic edits were made in Adobe Photoshop 2024 (R Core Team, 2020).

## 3.0 Results

### In silico *characterisation and allelic diversity of putative effector genes*

*EFC1, EFC2*, and *EFC3*, putative effector genes identified by Frampton & Shuey *et al*. (2024) are upregulated in the first 6 hours of infection by *A. psidii*, with transcript abundance significantly decreasing between 24 and 48 hours post infection. We identified alleles of these putative effector genes from the pandemic, South African, and Brazil (MF-1) genomes of *A. psidii*. We tested whether the encoded proteins meet the criteria of an effector using EffectorP; *EFC1* met the criteria of a cytoplasmic effector (Y-value=0.64), *EFC3* was predicted as both a cytoplasmic (Y-value=0.743) and an apoplastic effector (Y-value=0.504), and *EFC2* was not predicted to be a true effector (non-effector Y-value=0.594). We found one allele of *EFC2* in each of the genomes of *A. psidii*, which indicated that it is homozygous but there is a degree of allelic diversity across the population as each genome had a different allele. We identified at least two different alleles of *EFC1* and *EFC3* in available genomes of *A. psidii* variants, indicating these genes were heterozygous in the examined genomes. We located homologs of *EFC2* and *EFC1* in the Pucciniaceae, Phragmidiaceae, Sphaerophragmiaceae, Melampsoraceae, and Coleosporiaceae (Supplementary Figure 2), however, it is unclear if these homologs in related taxa are orthologous or paralogous.

### *dsRNAs targeting three putative* Austropuccinia psidii *effectors have variable impacts in vitro and in planta*

We tested the hypothesis that dsRNA-mediated inhibition of these putative effectors would supress *A. psidii* infection *in planta*, but not spore germination *in vitro*, as the effectors would be secreted from the infection architecture during host colonisation, after germination. We mixed urediniospores of *A. psidii* with dsRNAs specifically targeting each of these three effector genes and established *in vitro* spore germination assays as well as infection assays *in planta. In vitro*, dsRNA targeting *EFC2* almost completely inhibited germination (*p=0*.*0002*), as did the *EFC3* dsRNA (*p=0*.*007*), whereas dsRNA targeting variants of *EFC1* did not significantly inhibit spore germination compared to the –dsRNA control (*p=0*.*118*) or *GFP* non-specific dsRNA control (*p=0*.*645)* (Supplementary Figure 3). dsRNA treatments targeting different *EFC1* variants resulted in distinct differences in spore germination rates, which we hypothesised may be caused by differential activity or uptake of *EFC1* dsRNAs, further explained below. These results oppose our hypothesis that inhibition of putative effectors *in vitro* would have no effect and revealed that dsRNA the treatments inhibited germination, so that an infection could not occur after.

*In planta*, dsRNAs targeting *EFC1* variants (*p=0*.*022*) or *EFC2* (*p=0*.*019*) significantly inhibited infection and disease progression at two weeks post infection, with non-specific *GFP* dsRNA treatment having no effect (*p=0*.*779*) (Figures 1a, 1b). In a second *in planta* experiment, dsRNA targeting *EFC3* reduced disease symptoms at two weeks post-infection, but this was not significantly different to the controls (*p=0*.*072*) (Figure 1c, 1d). This result supported our hypothesis that inhibition of true effector genes would impact pathogen fitness and in turn, infection *in planta*. However, as shown in the *in vitro* experiments, inhibition of infection occurs through suppression of *A. psidii* spore germination, rather than through effector recognition by the host plant (Figure 1d).

**Fig. 1.**
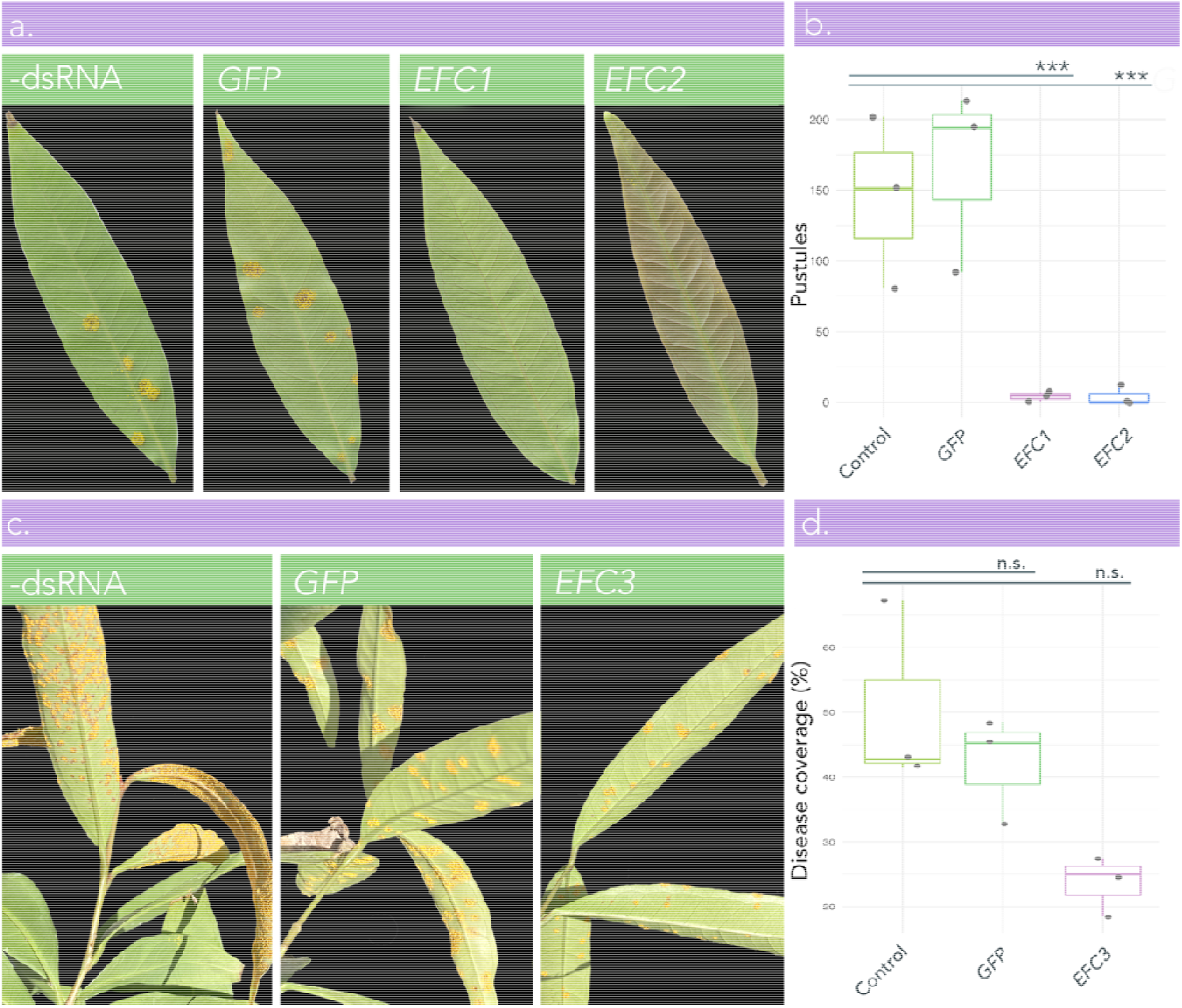
Impact of *Austropuccinina psidii*-specific dsRNA targeting two putative effectors on *Syzygium jambos*. *A. psidii* spores were co-inoculated with either nuclease-free H_2_O (–dsRNA negative control), *green fluorescent protein (GFP)* dsRNA (non-specific control), or dsRNA targeting putative effectors *EFC1, EFC2*, or *EFC3*. **(a)** Leaf scans of *S. jambos* leaves co-inoculated with *A. psidii* urediniospores and *A. psidii*-specific dsRNA (*EFC1* and *EFC2*), non-specific dsRNA (*GFP*), or H_2_O (–dsRNA control). Scans were taken at 2 weeks post-infection and each leaf is representative of other technical and biological replicates within its treatment group. **(b)** Boxplot with superimposed scatter (n=3) indicating the mean number of *A. psidii* pustules per leaf. Each biological replicate (plant) includes 4 technical replicates (leaves). Pustule counts were completed at 2 weeks post-inoculation. **(c)** Photographs of *S. jambos* plants co-inoculated with *A. psidii* urediniospores and *A. psidii*-specific dsRNA (*EFC3*), non-specific dsRNA (*GFP*), or H_2_O (–dsRNA control). Photographs were taken at 2 weeks post-infection. **(d)** Boxplot with superimposed scatter (n=3) indicating the disease coverage of *A. psidii* pustules on *S. jambos* leaves. Each biological replicate (plant) includes 4 technical replicates (leaves). Disease coverage assessments were completed at 2 weeks post-inoculation. Significance in boxplots (c), (d), and (f) is represented by asterisks (*=<0.05, **=<0.01, ***=<0.001 (Welch’s t-test)), n.s.= not significant. Bars in boxplots represent the standard error of the mean. Boxplots (a) and (b) were made in R 4.0.3.

### *Candidate effector EFC1 encompasses a gene-family of secreted proteins with high intragenomic diversity in* Austropuccinia psidii

We examined haplotype diversity of *EFC1* in the phased assembly of *A. psidii*. Three variants of putative effector *EFC1* (MK676_005453, MK676_005456/*EFC1-01* and MK676_005467/*EFC1-19*) clustered on chromosome 3A, occurring in tandem repeats flanked by transposable elements (Edwards et al., 2022), while a single variant (MK675_004127) was located on chromosome 3B (Figure 2a). The distance between these three closely linked orthologs on chromosome 3A is 11.40kb and 47.06kb respectively.

**Figure 2.**
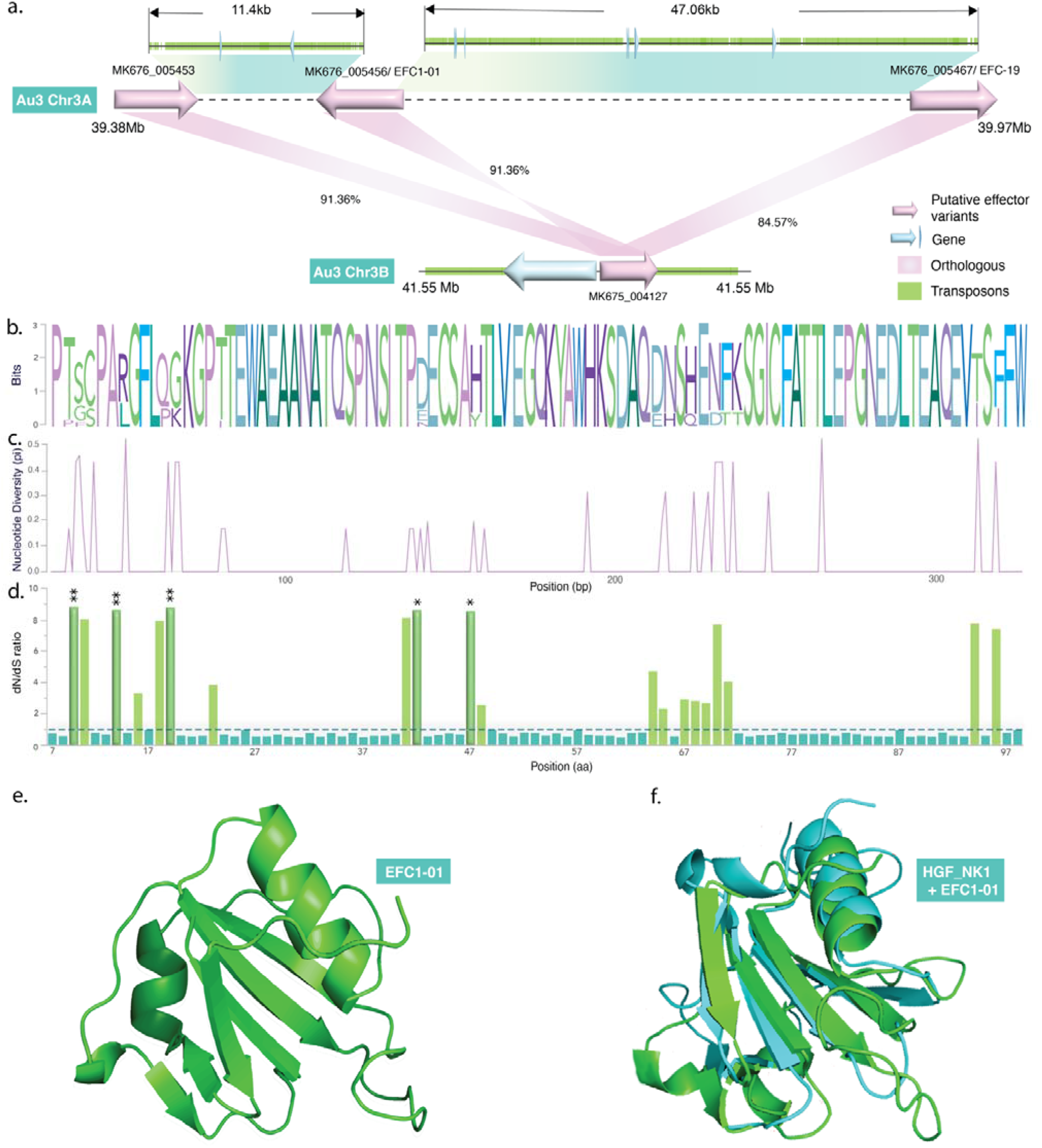
*In silico* characterisation of putative effector EFC1 and its variants in *Austropuccinia psidii*. (a) Genomic map of MK676_005453, MK676_005456 (*EFC1-01*), MK676_005467 (*EFC1-19*) and MK675_004127, putative effector gene variants in the EFC1 family (pink) in the *A. psidii* pandemic genome. *EFC1* variants are primarily clustered on Chromosome 3A and occur in tandem, flanked by transposable elements (green). (b) Sequence logo diagram illustrating the conservation of amino acids across 14 cloned, intragenomic variants of EFC1. (c) Nucleotide diversity (pi) graph showing sites of nucleotide polymorphisms across 14 variants of *EFC1*. (d) dN/dS plot illustrates rate of non-synonymous to synonymous substitution across 14 variants of EFC1. Asterisks indicated sites of statistically significant non-synonymous substitutions. Three sequences (EFC1-26, EFC1-28 and EFC1-06) with early terminations (stop codons) are excluded from the alignment used to produce sequence logo, pi plot, and dN/dS graphs. (e) Alphafold3 structural prediction of EFC1-01 protein. (f) Superimposition of EFC1-01 protein structure prediction and a Hepatocyte Growth Factor - NK1 fragment (HGF_NK1). Only the segment of the HGF_NK1 molecule that aligns with EFC1-1 is shown.

We amplified and cloned *EFC1* to obtain pure copies of variants identified in the annotated assembly and discovered sequence variants from cloned PCR product (Supplementary Figure 4, Supplementary 2). Two of these variants, *EFC1-01* and *EFC1-19* (*MK676_005456* and *MK676_005467*, respectively) occur in the assembled genome of *A. psidii*. We mapped RNA-seq reads to all variants of *EFC1* to determine whether they were transcribed (Supplementary dataset 1) and found evidence of overlapping reads with 100% identity mapping to majority of variants, except those with premature stop codons in their predicted translation (Supplementary Figure 4c). We determined the impact of SNPs of *EFC1* across all variants with measures of changes to translated proteins using a sequence logo (Figure 2b), nucleotide diversity (mean pi=0.0396, Figure 2c), and dN/dS ratio across the population of variants (Figure 2d). These measures of diversity indicate that variants of *EFC1* differ in their translated amino acids and may deviate due to non-neutral selection.

To gain insight into protein structure, function and potential binding interactions, we predicted the structure of *EFC-1* using Alphafold 3 (Abramson et al., 2024). The *EFC1-01* protein structure received a high predicted TM-score (PTM) of 0.81, and a Predicted Aligned Error (PAE) of 0.76, indicating a high degree of confidence for tertiary structure and relative positioning of residues within the protein, respectively (Figure 2e). The structure of *EFC1-01* is characterised by a beta-sheet core surrounded by alpha-helices. These features suggest a stable structure and loops and turns connecting secondary structure elements may allow for some flexibility and movement in the structure. The isoelectric point (pI) is 4.564 and the net charge at pH 7.4 is -8.3, indicating that the protein is acidic in nature and has a substantial number of acidic residues in comparison to basic, residues meaning that it may be more likely to interact with positively charged targets.

We compared our predicted structure with experimentally determined structures in the Protein Data Bank using the Dali server to determine homology with known proteins. (Holm, 2022). When superimposed, *EFC1-01* aligns closely with its best match in the PDB, the NK2 fragment of a Hepatocyte Growth Factor alpha chain (Figure 2f) (Holm, 2022). The structural alignment received a Z-score of 8.3, which is considered highly significant and infers that the structural similarities are non-random and that the molecules may share functional domains. The low (< 3 Å) root mean square deviation (RMSD) value of 2.5 Å (angstroms) indicates that the structures have overall similarity but may differ in minor features such as loop regions or side chains (Reva et al., 1998).

Together, the sequence logo, nucleotide diversity (pi), and dN/dS ratio plots illustrate the variable and conserved sites across putative variants of *EFC1* in the pandemic genome, including in the cloned copies. In most cases, nucleotide diversity corresponds with a non-synonymous amino acid substitution (Figures 2b, 2c, 2d). Variants of *EFC1* exhibit a higher rate of non-synonymous vs. synonymous substitutions (dN/dS ratio) compared to other coding sequences in the genome and the site model comparison indicates that the genomic variants of *EFC1* have evolved in a non-neutral manner in *A. psidii* (Figure 2d). The likelihood ratio test values for models M1a vs. M2a and M7 vs. M8 are all significant, indicating strong evidence for positive selection. The M2a and M8 models identify five codon sites (3, 7, 12, 35, and 40) out of 91 under positive selection (ω > 1 and *P* > 0.95).

### Two putative variants of EFC1 produce different phenotypes in vitro, but not in planta

We tested a hypothesis that the *EFC1* variants have different dsRNA inhibition phenotypes by conducting dsRNA-based *in vitro* and *in planta* assays using dsRNAs homologous to each of the *EFC1* variants. *EFC1-01* was the only treatment to not impact urediniospore germination, dsRNAs targeting all other variants of *EFC1* significantly reduced germination (Figure 3b). *EFC1-01* matched our expected inhibition pattern for an effector protein and was used for *in planta* assays. We included *EFC1-19* for comparison, as this was the only other cloned variant supported in the assembly of the pandemic genome.

**Fig. 3.**
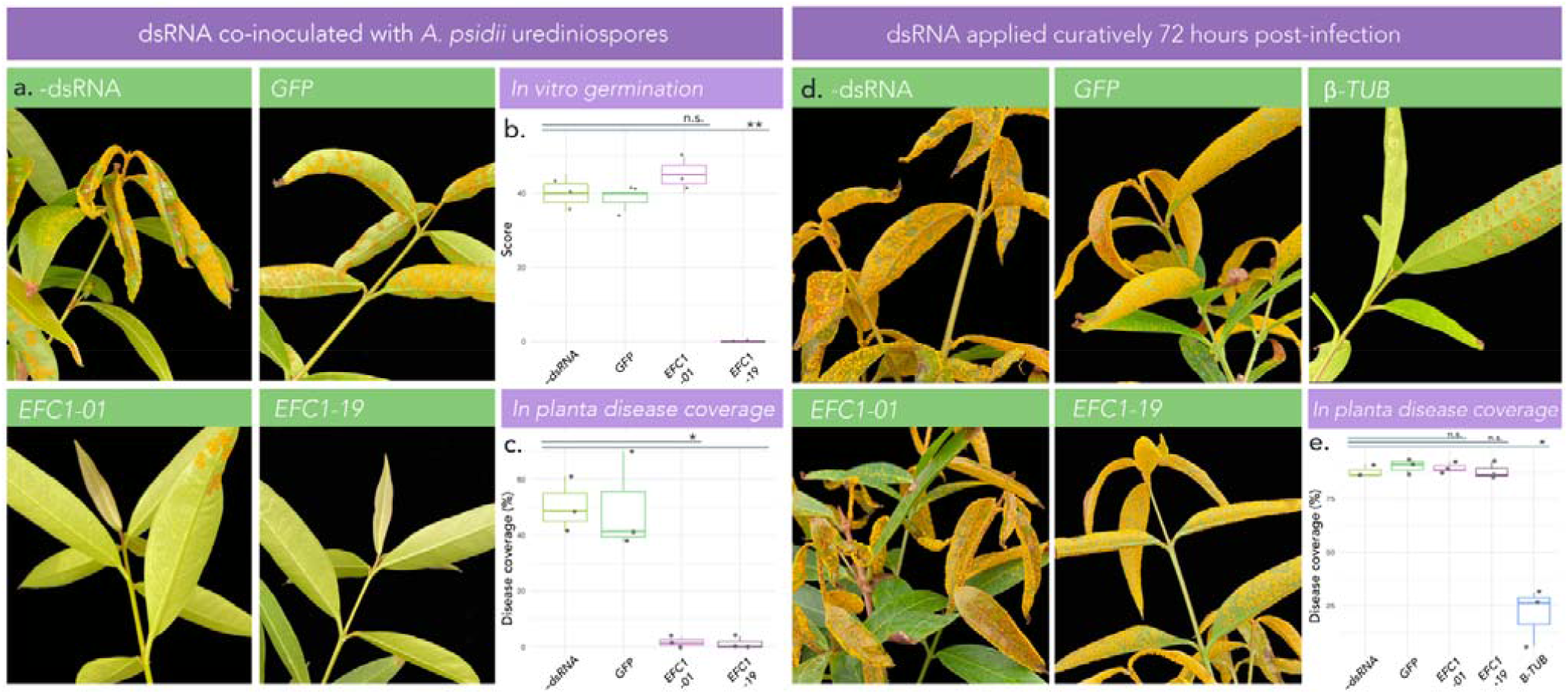
*Austropuccinia psidii*-specific dsRNA targeting two putative effector variants inhibits infection when co-applied with inoculum, but not when applied curatively. *A. psidii* spores were co-inoculated or curatively treated at 3 days post-inoculation with either nuclease-free H_2_O (–dsRNA negative control), green fluorescent protein (GFP) dsRNA (non-specific control), or dsRNA targeting closely-related variants (*EFC1-01* or *EFC1-19*) of putative effector *EFC1*. **(a)** Photo comparison shows plants co-inoculated with *A. psidii* urediniospores and *A. psidii*-specific dsRNA (*EFC1-01* or *EFC1-19*), non-specific dsRNA (*GFP*), or H_2_O (–dsRNA control). Photos were taken at 2 weeks post-inoculation. Background of photos was removed in Adobe Photoshop 2024. **(b)** Boxplot with superimposed scatter (n=3) showing percent (%) germination of *A. psidii* urediniospores *in vitro* on PVP-coated PDMS discs. Each biological replicate encompasses a count of at least 100 urediniospores to obtain the percent germination. Germination counts were completed at 24 hours post-inoculation. **(c)** Boxplot with superimposed scatter (n=3) indicating the mean percent (%) disease coverage of leaves. Each biological replicate (plant) includes 4 technical replicates that are the four youngest leaves on each plant. Pustule counts were completed at 2 weeks post-inoculation. **(d)** Photo comparison shows plants inoculated with *A. psidii* urediniospores and treated with *A. psidii*-specific dsRNA (*BTUB, EFC1-01* or *EFC1-19*), non-specific dsRNA (*GFP*), or H_2_O (–dsRNA control) at 3 days post-inoculation. Photos were taken at 2 weeks post-inoculation. Background of photos was removed in Photoshop (Beta). **(e)** Boxplot with superimposed scatter (n=3) indicating the mean percent (%) disease coverage of leaves. Each biological replicate (plant) includes 4 technical replicates that are the four youngest leaves on each plant. Pustule counts were completed at 2 weeks post-inoculation. Significance on the boxplot is represented by asterisks (*=<0.05, **=<0.01, ***=<0.001 (Welch’s t-test)). Bars represent the standard error of the mean. The boxplot was made in R 4.0.

*In vitro*, the co-inoculation of dsRNA targeting *EFC1* variant *EFC1-01* did not significantly reduce germination, as compared to the control (*p=0*.*2879*) and non-specific *GFP* dsRNA (Figure 3b). However, dsRNA targeting variant *EFC1-19* significantly inhibited spore germination (*p=0*.*0052*) (Figure 3b). We tested whether dsRNA of *EFC1-01* and *EFC1-19* would target each other by assessing sequence homology of all potential diced 21-mers starting at each base of the variant. Of 257 predicted 21-base pair fragments in dsRNA amplified for *EFC1-01* and *EFC1-19*, 104 were identical and 153 were non-identical (40% 21-mer homology) (Supplementary Figure 5). *In planta*, dsRNA treatments targeting variants *EFC1-01* and *EFC1-19* significantly inhibited disease coverage on *S. jambos* at 2 weeks post-infection (*p=0*.*0118* and *p=0*.*01104*, respectively). Non-specific *GFP* dsRNA controls did not inhibit disease coverage *in planta*, as compared to the –dsRNA control (*p=0*.*948*) (Figures 3a, 3c).

### Curative dsRNA targeting putative effector EFC1 has no impact on myrtle rust infection when applied at 3 days post-inoculation

Following *in vitro* and *in planta* co-inoculation assays, putative effector variant *EFC1-01* appeared to align with expected effector phenotypes; germination was not inhibited *in vitro* but was inhibited *in planta* (Figures 3a, 3b, 3c). Time course transcript data from Frampton & Shuey *et al*. (2024) suggests that *EFC1* expression decreases significantly between 24- and 48-hours post-inoculation. Based on this knowledge, we tested a hypothesis that curative dsRNA treatments targeting *EFC1-01* at 72 hours post-infection would not significantly decrease myrtle rust symptoms *in planta* at 2 weeks post-infection.

Curatively applied dsRNA targeting putative effector variants *EFC1-01* and *EFC1-19* did not decrease disease coverage *in planta* (*p=0*.*4993* and *p=1*, respectively). Non-specific *GFP* dsRNA controls did not inhibit disease coverage *in planta*, as compared to the –dsRNA control (*p=0*.*4204*). Plants treated with *BTUB* dsRNA had significantly reduced disease coverage (*p=0*.*0102*). A positive control (*BTUB*) was included to ensure that curatively applied dsRNA could effectively reduce disease coverage *in planta*, as has been previously shown (Degnan et al., 2023b) (Figures 3d, 3e).

## 4.0 Discussion

Specialist and generalist pathogens evade recognition by manipulating the host response through a suite of secreted proteins called effectors. We examined putative effector genes in *Austropuccinia psidii*, an invasive rust fungus with a wide host rangeto characterise their roles in infection before haustorial development. Our co-inoculated and curative dsRNA treatments showed these putative effectors are crucial to initiate but not maintain an infection (Figures 3a–3e). Inhibition of two variants belonging to a putative effector with high intragenomic variability (*EFC1*) caused differential symptoms *in vitro*. Significant rates of non-synonymous mutation were observed across variants (Figure 2d), akin to non-neutral selection or adaptive evolution, which may indicate that the expression profile of this gene family benefits from having multiple functional variants that potentially facilitate pathogenicity across the almost 500 species of Myrtaceae currently known to be infected by *A. psidii*.

*Austropuccinia psidii* may benefit from having several functionally different variants of *EFC1* that are expressed at different rates, and variation at this putative effector may be a strategy to outpace evolution of disease recognition or to allow for flexibility in the infection of the almost 500 known hosts. A single variant of *EFC1-19* had a significantly higher average RNA-seq read depth (suggesting higher expression) than other variants (Supplementary Table 2), and two variants were likely pseudogenes based on premature stop codons. The ability to infect a wide range of hosts or evade recognition can be further realised through non-neutral selection-driven diversification of proteins, and perhaps amplified through transposable element (TE) insertion events (Lo Presti et al., 2015; Raffaele and Kamoun, 2012). Alternatively, many fungal pathogens also exhibit redundancy in effector genes; variants of *EFC1* may indicate a suite of effectors that work together to facilitate infection, providing a backup mechanism to counteract host defences if other effectors are neutralised (Win et al., 2012).

The intragenomic variability observed in *EFC1* may advantage clonal populations if urediniospores express genomic variants that are functionally different. The outcome of built-in genetic diversity has the advantage of testing variants without the need for evolutionary innovation through recombination. Intragenomic variation of a putative effector in *A. psidii* has similarities to the mode-of-infection used by the malaria parasite, *Plasmodium falciparum*, which switches expression of *var* genes, a large multi-gene family that contributes to the pathogens escape from recognition by the host immune system by altering surface antigens (Goel et al., 2010). Switches in expression of v*ar* genes are often regulated through histone modifications, such as acetylation and methylation (Goel et al., 2010). These modifications are maintained by mutually exclusive repression, a state in which only one (*var*) gene is expressed while others are kept in a silent state (Mok et al., 2008). If *A. psidii* has converged on a similar strategy for pathogenicity, repression may explain why one variant is expressed more so than others. In both *P. falciparum* and hypothetically, *A. psidii*, this infection strategy has two benefits: quick adaptation to gene-for-gene resistance, and opportunities to trial many functionally different effectors from one genotype that produces millions of spores.

There was evidence for four genomic variants of *EFC1* in the assembly of *A. psidii*, and our discovery of multiple cloned variants was unexpected. An alternative hypothesis to the observed intragenomic diversity of *EFC1* is that the variants are an artefact of PCR, whether due to copy error or to PCR recombination, and not a true reflection of diversity in the genome. We used a high-fidelity polymerase, cloned each variant, and supported most variants with reads from RNA-seq, thus propose that PCR-error is a less parsimonious explanation than intragenomic variation as several independent SNP errors in early rounds of amplification would be needed to generate 13 variant sequences in the PCR product. PCR recombination may explain two of the variants, *EFC1-19* and *EFC1-33*, based on the SplitsTree analysis, however, *EFC1-19* is present in the genome assembly, and recombination at these complex loci may be an outcome of sexual reproduction. We amplified and cloned only two of four variants detected in the assembled genome of *A. psidii*, and our data therefore suggests that more *EFC1* variants may exist in the genome. We show that these variants that differ by a few SNPs occur in tandem repeats flanked by TEs, a phenomenon with precedence in the smut fungi (Dutheil et al., 2016). These regions will be difficult to assemble, unless sequenced across the entire repeat.

*EFC1* is expressed before haustorial formation and is crucial for infection; knowledge of its function may be key to understanding the initial stages of pathogen infiltration and disease evasion (Ultsch et al., 1998). We modelled EFC1 and found structural similarities between EFC1-01 (PTM=0.81) and the NK1 heparin-binding domain complex of a hepatocyte growth factor (NK1-HGF) (Z-Score=8.3). The N-terminal hairpin heparin-binding domain of NK1-HGF gives the structure intrinsic stability and suggests a simple ancestral protein with a conserved motif (Zhou et al., 1998). Based on the high Z-score in the structural alignment of EFC1-01 and NK1-HGF, indicating these proteins may share similarities in function, and the knowledge that NK1-HGF binds heparin gylcosaminoglycans (GAGs), we hypothesise that putative effector EFC1-01 binds structurally similar polysaccharides to facilitate infection in the *A. psidii* pathosystem GAGs are long, unbranched polysaccharides, and share structural similarity with plant-cell wall pectins (Etzler and Mohnen, 2009), suggesting that a possible function of EFC1-01 is the binding of pectin (or other plant cell wall polysaccharides) to facilitate infection by *A. psidii*. This putative function is demonstrated in *Pseudocercospora fuligena*, where binding of de-esterified pectin by *Pf*Avr4-2 interferes with Ca^2+^-mediated cross-linking, to loosen plant cell structure and impede its integrity (Chen et al., 2021). This is an abnormal behaviour for an Avr4-family protein, as effectors of this family typically bind chitin (Van Den Burg et al., 2006). Functional assays, such as immunolocalization or fluorescent labelling assays could be used to further explore this hypothesis (Chen et al., 2021).

Extensive flanking of putative effector *EFC1* variants by TEs suggests that TEs may have facilitated duplication of new variants (Figure 2a). Such outcomes may arise through TE copy/paste insertion events, or non-allelic homologous recombination, which can occur in the presence of multiple TEs (Feschotte and Pritham, 2007; Kidwell and Lisch, 2001). Effector proteins evolve quickly in a co-evolutionary arms race with host defence mechanisms (Jones and Dangl, 2006). TEs further facilitate this rapid diversification of effectors and are often seen as the key drivers of rapid pathogen adaptation (Fouché et al., 2018). The high homology of *EFC1* variants on Chr3A and Chr3b suggests that this putative effector family has duplicated or diverged while conserving core functions (Figure 2a). *A. psidii* has an uncharacteristically broad host range, thus the TE-facilitated diversification of secreted proteins crucial for early-stage infection may be linked to its ability to infect and overcome plant defences in a wide range of Myrtaceous hosts. In addition, homologs of *EFC1* were not multi-allelic in other members of the Pucciniales and we could not determine if they were orthologs or paralogs (Supplementary Figure 2). Future studies could clone and functionally test this effector to determine if intragenomic variation is observed in other taxa, or if the proliferation of this putative effector is confined to *A. psidii*.

Given that effectors are secreted during colonisation of the host plant, we hypothesised that dsRNA-based inhibition of putative effectors would prevent symptoms *in planta* but have no effect *in vitro* without host contact (Petre et al., 2014; Rafiqi et al., 2010). dsRNA targeting *EFC1-01* partially inhibited spore germination while dsRNA targeting *EFC1-19* fully inhibited germination with *in vitro* (Supplementary Figure 3). This result highlights gaps in our understanding of exogenous dsRNA-mediated inhibition in fungi. We showed that 40% of 21-mers from dsRNA targeting *EFC1-01* would be homologous with *EFC1-19*, however this was not enough to cause inhibition (Supplementary Figure 5). We do not have a definitive explanation for this outcome, but it may be linked to the difference in expression between the two tested variants (*EFC1-19* is more expressed than *EFC1-01*) or that 40% sequence homology is insufficient to inhibit a non-targeted variant.

Time course RNA-seq data reveals that *EFC1* expression decreases drastically between 24–48 hours (Frampton et al., 2024). In response, we hypothesised that 3 days post-infection curative applications of dsRNA targeting *EFC1-01* and *EFC1-19* would have no impact on disease *in planta*. Curative assays, which included *BTUB* dsRNA treatments as a positive control, supported this hypothesis showing no changes in disease coverage in *EFC1* dsRNA-treated plants (Figures 3d, 3e). Together, the results support the hypothesis that *EFC1* is crucial for onset, but not maintenance of infection in the *A. psidii* pathosystem, and its inhibition phenotypes fulfill our expectations of an effector protein. Given their early expression in infection, and inhibition phenotypes observed *in vitro* and *in planta*, both *EFC2* and *EFC3* are likely genes associated with germination, or infection expressed by germinating urediniospores, early in the infection process, and based on Effector P analysis, *EFC3* is predicted as a cytoplasmic and apoplasmic effector, but further work is needed to confirm their roles in infection.

Although we did not confirm how *EFC1* interacts with its host at the molecular level, several lines of evidence support this gene being an effector. Our findings suggest that the *EFC1* gene family plays a significant role in early-stage infection by *A. psidii*, with dsRNA-based inhibition indicating that two variants are crucial for initiation, but not maintenance of infection. Intragenomic variability within this gene family, along with a signature of non-neutral selection, leads to the hypothesis that diversification of *EFC1* variants could be driven by adaptive evolution and further enhanced through TE-facilitated evolution, contributing to an uncharacteristically broad host range of *A. psidii*. Structural similarities between *EFC1-01* and NK1-HGF hint at a potential role in cell-wall binding, and further support the hypothesised role of *EFC1* as an effector. Finally, at its centre of origin in Brazil, *A. psidii* is a minor pathogen of native Myrtaceae. Developing a better understanding of pathogenicity and resistance in native pathosystems may help resolve the role that effectors play in assisting invasive pathogens such as *A. psidii* to establish such a broad host range.

## CRediT author statement

**Rebecca Degnan:** conceptualisation, methodology, software, validation, formal analysis, investigation, data curation, writing – original draft, writing – review and editing, visualisation, project administration. **Anne Sawyer:** conceptualisation, methodology, resources, writing – review and editing, supervision, project administration, funding acquisition. **Donald Gardiner:** conceptualisation, methodology, writing – review and editing, supervision. **Zhenyan Luo:** software, formal analysis, investigation, data curation, writing – review and editing, visualisation. **Rebekah Frampton:** conceptualisation, resources, writing – review and editing. **Benjamin Schwessinger:** resources, data curation, writing – review and editing, supervision. **Neena Mitter:** resources, data curation, writing – review and editing, supervision. **Bernard Carroll:** conceptualisation, methodology, resources, writing – review and editing, supervision, funding acquisition. **Grant Smith:** conceptualisation, methodology, resources, writing – review and editing, supervision, funding acquisition. **Alistair McTaggart:** conceptualisation, methodology, software, validation, formal analysis, data curation, writing – original draft, writing – review and editing, visualisation, supervision, funding acquisition, project administration, **Louise Shuey:** conceptualisation, methodology. resources, writing – review and editing, supervision, funding acquisition.

## Acknowledgments

We thank the Australian Plant Biosecurity Foundation (APBSF) for their support. R.M.D thanks the Plant Biosecurity Research Initiative (PBRI). A.S was supported by an Advance Queensland Industry Research Fellowship (AQIRF095-2021RD4). This project was partially funded with a grant from the New Zealand Ministry of Business, Innovation & Employment (MBIE) Endeavor Fund, Beyond Myrtle Rust project (#C09×1806).

## Data sharing statement

Read mapping, coding sequences of all variants, effector candidates, and homologs in other rust fungi, as well as raw *in vitro* and *in planta* data that support the findings of this study are made publicly available through Zenodo (10.5281/zenodo.14238932). Any additional data can be made available upon request.

---

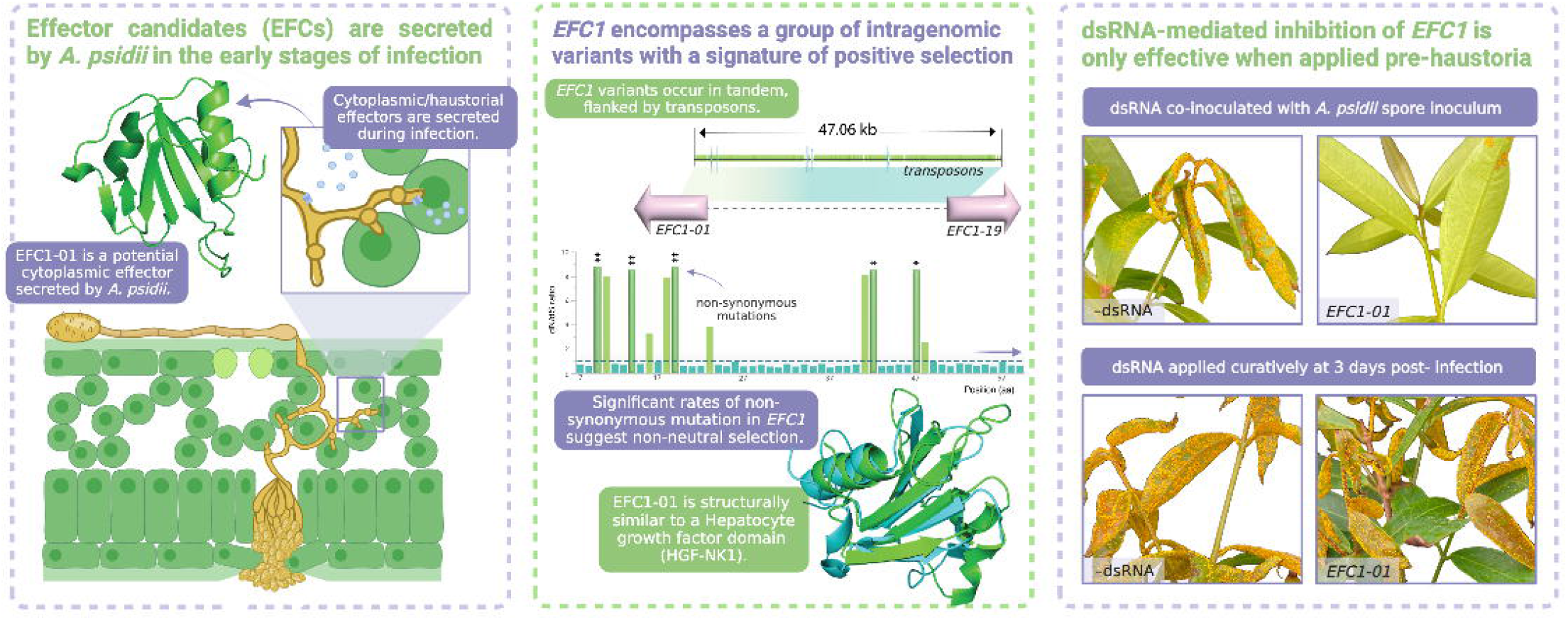

