## Supplementary file for "Intragenomic variants of a putative effector drive early-stage infection in a broad host-range rust fungus"

**Supplementary Table 1. Primer sequences for amplification of Effector Candidates 1, 2, and 3 (EFC1/2/3).** All primers were also ordered with the T7 sequence (TAATACGACTCACTATAGGG) at the 5’ end to allow for synthesis of double stranded RNA. Amplicon length (bp) does not include introns.

| **Effector** | **Direction** | **Primer sequence (5’-3’)** | **TM (ºC)** | **Amplicon length (bp)** | **No. introns** | **Total length introns (bp)** |
| --- | --- | --- | --- | --- | --- | --- |
| EFC1 | Forward | *CTTTTAGGCTTGATGCTTG* | 58.2 | 322 | 5 | 386 |
|  | Reverse | *GTCATTCATAGGGATCATG* | 55.6 |  |  |  |
| EFC2 | Forward | *GTGCTCTCTACTTTACTCC* | 51.0 | 436 | 1 | 95 |
|  | Reverse | *GAGGTTCATCTGTAATTG* | 50.4 |  |  |  |
| EFC3 | Forward | *CTTTGGCTCAAACAATGAC* | 58.8 | 357 | 2 | 290 |
|  | Reverse | *GATTCGGTAGGTCTGATG* | 55.9 |  |  |  |

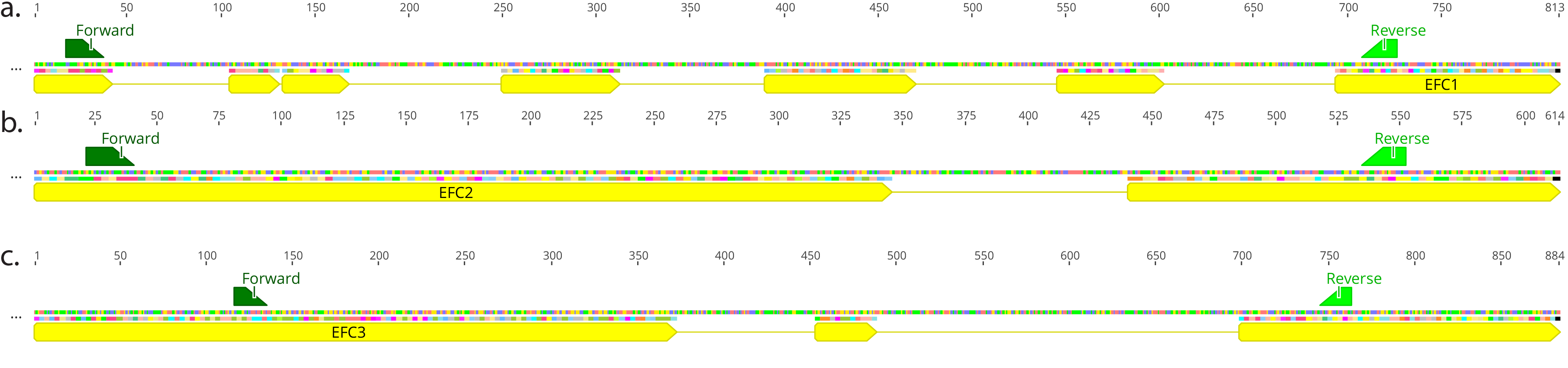
**Supplementary Figure 1. Primer placements for design of dsRNA targeting putative effectors in *Austropuccinia psidii*.** Yellow annotations indicate coding sequences and green annotations indicated forward and reverse (labelled) primers for **(a)** EFC1, **(b)** EFC2, and **(c)** EFC3.

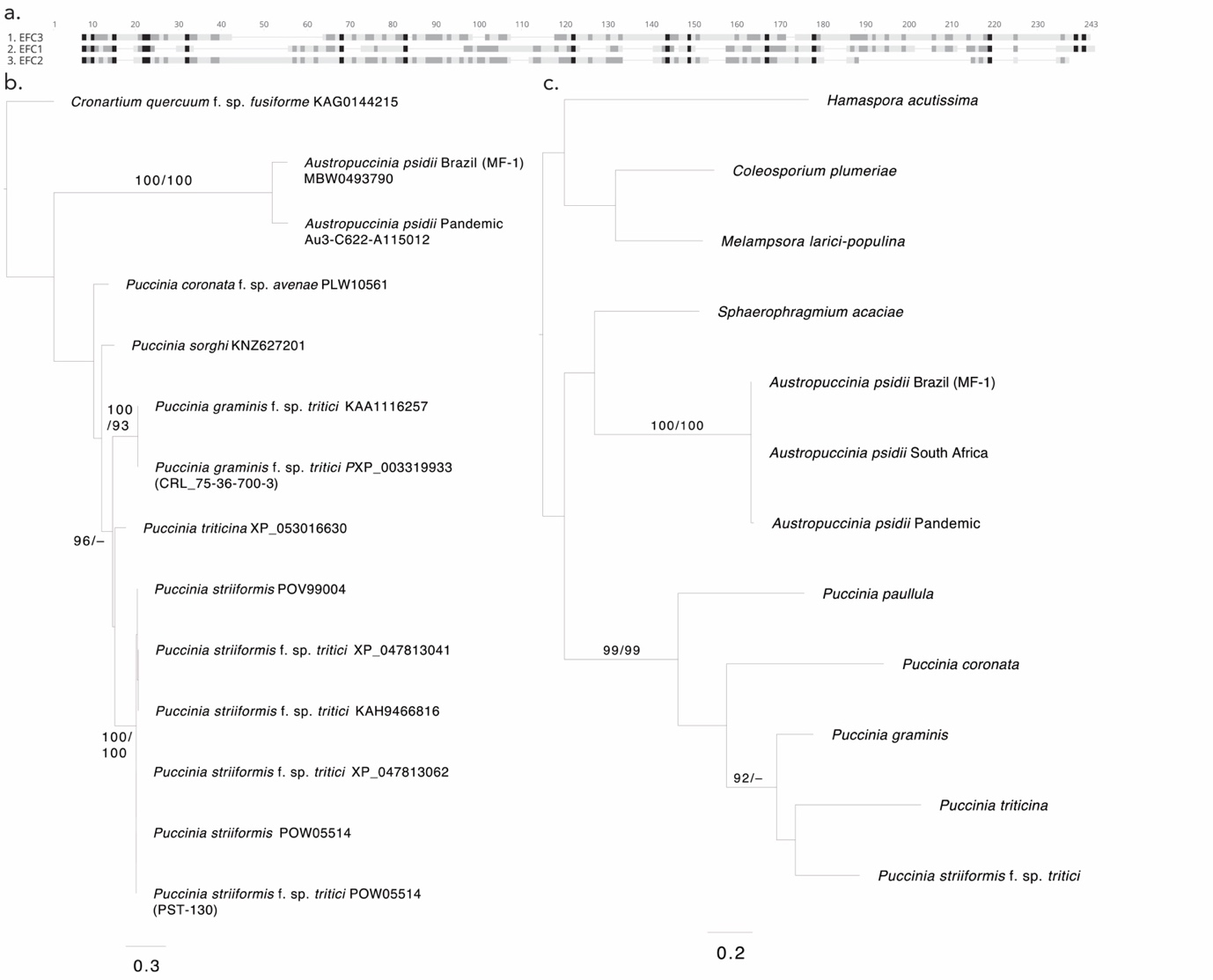

**Supplementary Figure 2. Homology of Effector Candidates 1, 2, and 3 (EFC1/2/3).** (a) Amino acid alignment of EFC1,2, and 3. (b) Phylogenetic tree of EFC1 homologs in related taxa. (c) Phylogenetic tree of EFC2 homologs in related taxa.

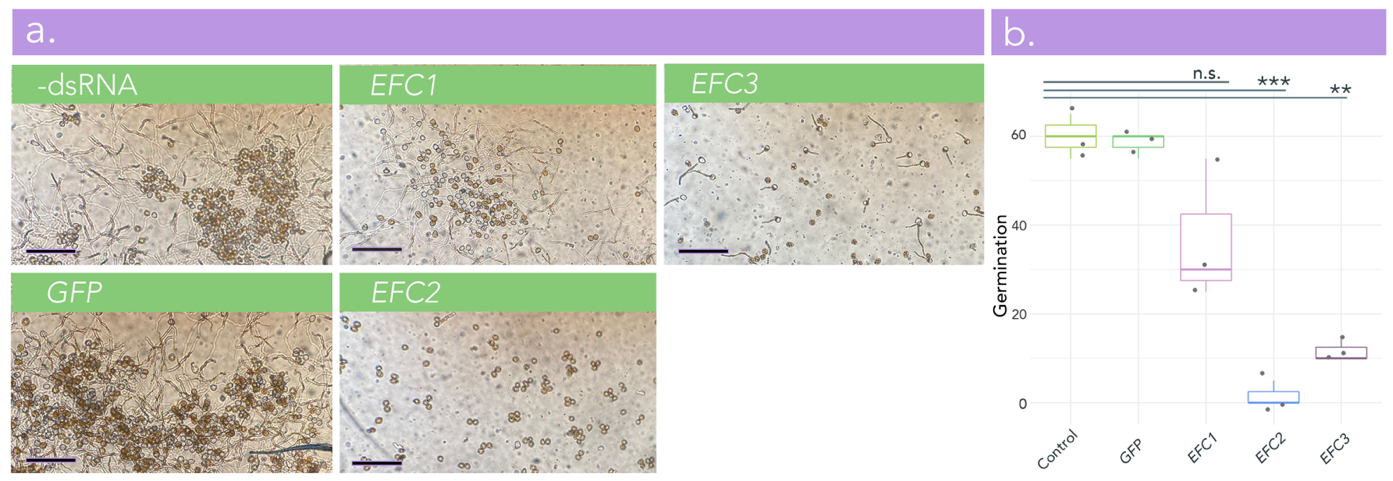
**Supplementary Figure 3. In vitro germination of *Austropuccinia psidii* urediniospores treated with dsRNA targeting three effector candidates (EFCs). (a)** Photographs of control and dsRNA-treated *A. psidii* urediniospores on PVP-coated PDMS discs at 24 hours post-inoculation. Photographs taken at 40x10 magnification using the EP view application. **(b)** Boxplot with superimposed scatter (n=3) showing percent (%) germination of *A. psidii* urediniospores *in vitro* on PVP-coated PDMS discs. Each biological replicate encompasses a count of at least 100 urediniospores to obtain the percent germination. Germination counts were completed at 24 hours post-inoculation. Germination scores taken at 24 hours post infection. Significance on the boxplot is represented by asterisks (*=<0.05, **=<0.01, ***=<0.001 (Welch’s t-test)). Bars represent the standard error of the mean. The boxplot was made in R 4.0.3.

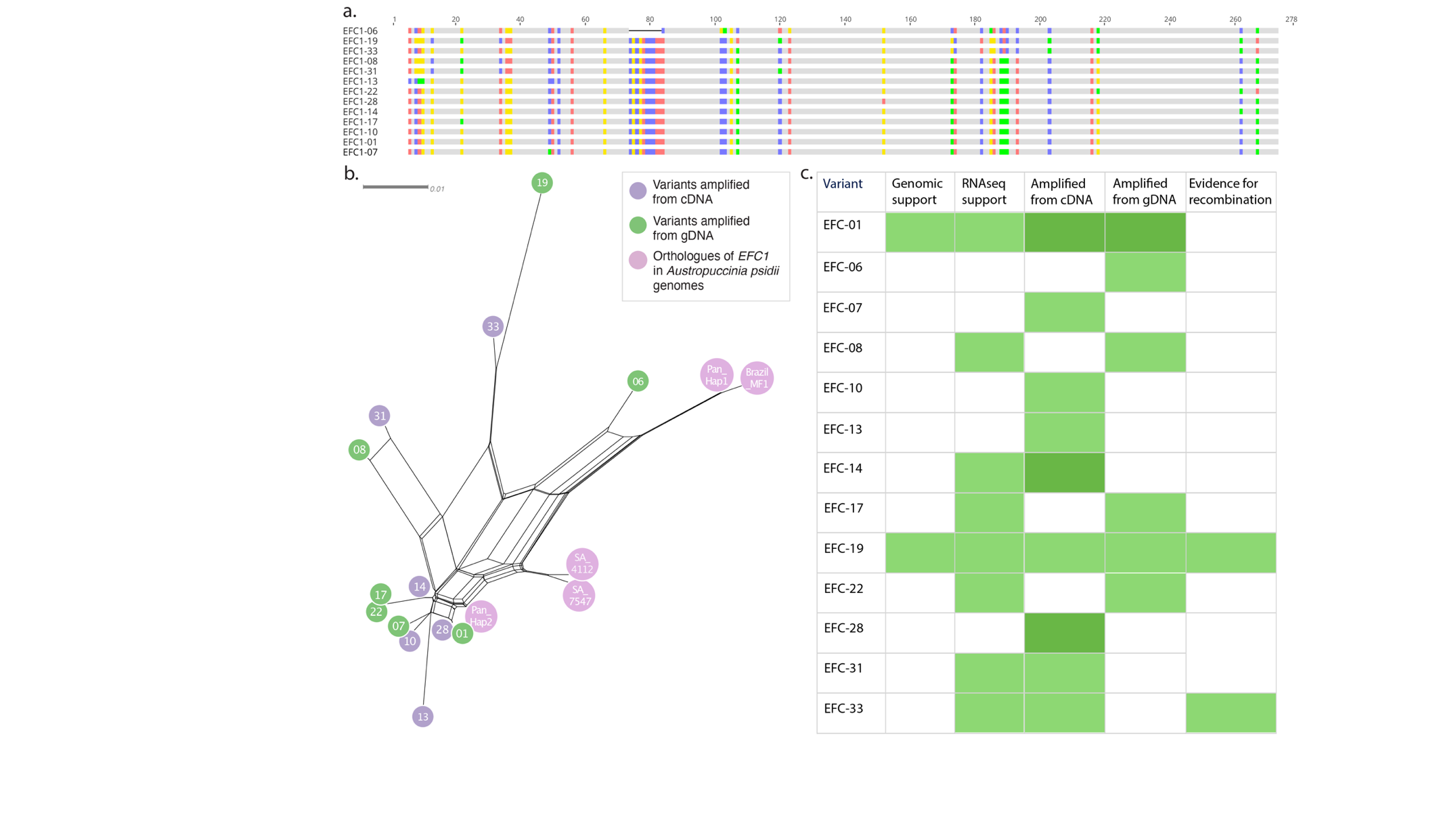

**Supplementary Figure 4. Sequence homology, phylogenetics, and summary of support for Effector Candidate 1 (EFC1) putative variants in *Austropuccinia psidii.*** (a) Clustal omega (similarity organised) alignment of *EFC1* putative variants cloned from *A. psidii.* Sequence mismatches are highlighted. (b) Phylogenetic tree of *EFC1* putative variants and orthologues in available *A. psidii* genomes, produced with MAFFT alignment and visualised using SplitsTree. Purple circles = EFC1 variants amplified from cDNA; green circles = *EFC1* variants amplified from gDNA, pink circles = orthologues of *EFC1* in available genomes. (c) Table summarising support for *EFC1* putative variants: no shading = no support; light green shading = support; dark green shading = variant was captured in sampling (cloning, colony selection, amplification, and sequencing), more than once. Genomic support based on most recent *A. psidii* genome assembly (APSI_hap1_v2, GCA_023105745.1). RNAseq support based on mapping with 100% sequence homology. Evidence for recombination based on significant result in Chi-squared test.

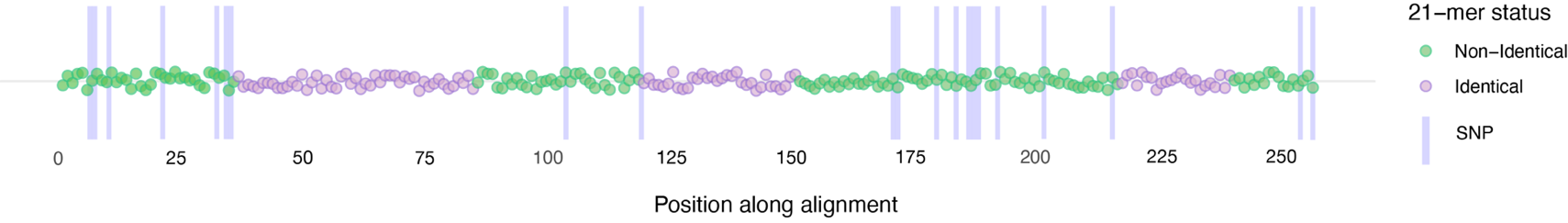
**Supplementary Figure 5. Raindrop plot of 21-mer status in EFC1-01 versus EFC1-19 in relation to single nucleotide polymorphisms (SNPs) and position along alignments.** Dots represent starting nucleotide of the 21-mer. Green = non-identical 21-mers; pink = identical 21-mers, purple bars = SNPs.

**Supplementary Table 2.** Sequence variation and average read depth of the *Effector Candidate 1* (*EFC1*) variants in *A. psidii*. All EFC1 protein sequences are 91 amino acids in length. Read mapping is with 100% sequence identity. *EFC1-28* and *EFC1-06* include premature stop codons.

| Putative *EFC* gene variant | % amino acid identity with *EFC1-01* | % nucleotide identity with *EFC1-01* | Number of non-synonymous mutations (compared to *EFC1-01*) | Mean read depth at 24 hours post-inoculation |
| --- | --- | --- | --- | --- |
| *EFC1-01* | 100 | 100 | 0 | 233.9 |
| *EFC1-06* | 30.2 | 91.6 | 61 | 223.4 |
| *EFC1-07* | 98.9 | 99.6 | 1 | 206.3 |
| *EFC1-08* | 94.5 | 97.5 | 5 | 1332.1 |
| *EFC1-10* | 98.9 | 99.6 | 1 | 149.9 |
| *EFC1-13* | 97.8 | 98.6 | 2 | 149.9 |
| *EFC1-14* | 98.9 | 99.6 | 1 | 221.1 |
| *EFC1-17* | 98.9 | 99.6 | 1 | 220.4 |
| *EFC1-19* | 89.0 | 92.4 | 14 | 4703.6 |
| *EFC1-22* | 97.8 | 98.9 | 2 | 391.3 |
| *EFC1-28* | 98.9 | 99.6 | 1 | 234.6 |
| *EFC1-31* | 92.3 | 96.8 | 7 | 1232.91 |
| *EFC1-33* | 91.3 | 95.7 | 7 | 842.1 |
